# Melanin-concentrating hormone promotes feeding through the lateral septum

**DOI:** 10.1101/2024.05.20.595067

**Authors:** Mikayla A. Payant, Anjali Shankhatheertha, Melissa J. Chee

## Abstract

Feeding is necessary for survival but can be hindered by anxiety or fear, thus neural systems that can regulate anxiety states are key to elucidating the expression of food-related behaviors. Melanin-concentrating hormone (MCH) is a neuropeptide produced in the lateral hypothalamus that promotes feeding and anxiogenesis. The orexigenic actions of MCH that prolong ongoing homeostatic or hedonic feeding are context-dependent and more prominent in male than female rodents, but it is not clear where MCH acts to initiate feeding. The lateral septum (LS) promotes feeding and suppresses anxiogenesis when inhibited, and it comprises the densest projections from MCH neurons. However, it is not known whether the LS is a major contributor to MCH-mediated feeding. As MCH inhibits LS cells by MCH receptor (MCHR1) activation, MCH may promote feeding via the LS. We bilaterally infused MCH into the LS and found that MCH elicited a rapid and long-lasting increase in the consumption of standard chow and a palatable, high sugar diet in male and female mice; these MCH effects were blocked by the co-administration of a MCHR1 antagonist TC-MCH 7c. Interestingly, the orexigenic effect of MCH was abolished in a novel, anxiogenic environment even when presented with a food reward, but MCH did not induce anxiety-like behaviors. These findings indicated the LS as a novel region underlying orexigenic MCH actions, which stimulated and enhanced feeding in both sexes in a context -dependent manner that was most prominent in the homecage.

## 1. Introduction

Melanin-concentrating hormone (MCH) is a hypothalamic neuropeptide known to regulate several behaviours including feeding, the first ascribed role of MCH (Qu et al., 1996). *Mch* mRNA expression is elevated in obesity, and this increase is linked with hunger, as fasting also promotes *Mch* expression (Qu et al., 1996). These transcriptional changes suggested that MCH promotes feeding, and indeed, MCH administration into the lateral (Qu et al., 1996; Kela et al., 2003) or third ventricles (Rossi et al., 1997; Duncan et al., 2005) acutely increased chow intake and sustained this orexigenic effect for up to six hours (Qu et al., 1996; Rossi et al., 1997; Kela et al., 2003). Moreover, the chronic intake of palatable high-fat diets also elevated the gene expression for *Mch* and the MCH receptor, *Mchr1* (Elliott et al., 2004). By contrast, *Mch* deletion (Pissios et al., 2008; Mul et al., 2011) or MCHR1 antagonism (Karlsson et al., 2012) attenuated the intrinsic reward properties of palatable food.

MCH has been associated with multiple aspects of feeding behaviour but is most consistently involved in prolonging food bouts to promote overeating (Lee et al., 2021). MCH neurons are active during, but not preceding, food consumption (Subramanian et al., 2023), and while activating MCH neurons did not consistently stimulate feeding, activating MCH neurons during a food bout increased the amount of food consumed (Dilsiz et al., 2020). Consistently, MCHR1 deletion prevents cue-induced overeating (Sherwood et al., 2015), which may be related to a role for MCH in improving taste evaluation and taste-independent reward associated with nutrient-dense foods. MCH infusion increased the initial rate of licking a sweet, but not a bitter, solution suggesting that increased intake was based on positive taste (Baird et al., 2006). MCHR1 antagonism also reduced effort for a sucrose reward but not a saccharin reward (Karlsson et al., 2012), so MCH preferentially reinforces calorically-dense substances. Consistent with the role of MCH to prolong food bouts or extend ongoing feeding, the orexigenic effects of MCH were more robust in the dark than light cycle when rodents have already initiated feeding (Terrill et al., 2020).

Along with promoting food intake, MCH also has a role in anxiety-like behaviour and therefore may regulate context-dependent, anxiety-related food avoidance. MCH administration can both increase (Smith et al., 2006) and decrease anxiety (Kela et al., 2003), therefore it may have synergistic or opposing roles on promoting food intake in an anxiogenic environment. In addition, given the involvement of MCH neurons in memory (Izawa et al., 2019; Liu et al., 2022), it is plausible that MCH action at downstream sites could help integrate contextual information to determine when it is appropriate to prolong eating.

The distributions of MCH projections and MCHR1 are widespread throughout the brain, but the brain regions underlying MCH-mediated feeding are not fully characterized. The orexigenic effects of MCH have largely been described in rats by infusing MCH into discrete brain regions, which included the arcuate nucleus (Abbott et al., 2003), paraventricular nucleus (Rossi et al., 1999; Abbott et al., 2003), and lateral hypothalamic area (Romero-Pico et al., 2018). Notably, the orexigenic effects of MCH are most prominently featured in the nucleus accumbens (Georgescu et al., 2005; Guesdon et al., 2009; Terrill et al., 2020), where MCH infusion promoted both homeostatic and hedonic feeding (Terrill et al., 2020). However, the orexigenic effects of MCH have only been reported in male subjects (Terrill et al., 2020), as MCH-mediated feeding is inhibited by estradiol in female subjects (Messina et al., 2006; Santollo and Eckel, 2008). While MCH infusion in the nucleus accumbens consistently promoted feeding, activating accumbens-projecting MCH neurons has yielded mixed results (Noble et al., 2018; Terrill et al., 2020). Chemogenetic activation of MCH neurons projecting to the nucleus accumbens increased chow intake in both the light and dark cycle (Terrill et al., 2020), however, in a different study, the same procedure had no effect on chow intake in the light cycle (Noble et al., 2018).

Given the relative paucity of targets supporting the orexigenic actions of MCH, we determined if the lateral septum (LS) is a target site of MCH action. The LS, especially along its lateral border (Payant et al., 2023), receives dense projections from MCH neurons (Chee et al., 2015) and expresses MCHR1 (Lembo et al., 1999; Saito et al., 2001; Chee et al., 2013; Diniz et al., 2020), and MCH can directly inhibit LS cells (Liu et al., 2022; Payant et al., 2023). LS inhibition stimulates feeding (Mitra et al., 2014; Gabriella et al., 2022), thus MCH-mediated inhibition in the LS may promote food intake. We infused MCH bilaterally into the LS of male and female wildtype mice and determined homecage feeding of regular chow or a highly palatable sugar pellet. Since the LS can regulate both feeding and anxiety-like behaviour (Bakshi et al., 2007; Xu et al., 2019), we also assessed whether food availability promoted entry into anxiogenic environments. We found that intra-LS MCH infusion significantly increased the consumption of chow or a palatable diet in both males and females, especially when MCH was infused directly above MCHR1-rich LS regions (Payant et al., 2023). However, when a food reward like a familiar palatable food pellet was placed in the center of an open field, which engenders avoidant anxiety-like behaviours, MCH did not facilitate food exploration in sated or fasted mice. Taken together, these findings highlight the LS as an important structure underlying MCH-induced feeding, but this orexigenic effect was present only in the habituated homecage and not apparent in novel, anxiogenic environments.

## 2. Materials and Methods

All procedures described herein were approved by the Carleton University Animal Care Committee in accordance with guidelines provided by the Canadian Council on Animal Care. All C57BL/6J wildtype mice (stock 000664; Jackson Laboratory, Bar Harbor, ME) were bred in-house, maintained on a 12-hour light-dark cycle (22–24 °C; 40–60% humidity), and given *ad libitum* access to regular chow (2.9 kcal/g; Teklad Global Diets 2014, Envigo, Mississauga, Canada) or 60% high dextrose diet (3.6 kcal/g; Teklad Custom Diet, TD.05256, Envigo, Mississauga, Canada) and water.

### 2.1 Cannula implantation

Male and female mice (8–14 weeks old) received a subcutaneous (sc) injection of carprofen (20 mg/kg; Zoetis, Kirkland, QC, Canada) analgesia 30 minutes prior to the start of surgery. They were then anesthetized with isoflurane, secured in a stereotaxic apparatus (David Kopf Instruments, Tujunga, CA), and their incision sites were sterilized before making a rostrocaudal incision to expose the skull. The skull surface was cleaned and air-dried, and the head position was leveled within the stereotaxic frame using bregma and lambda coordinates read from the skull.

A bilateral 26 gauge stainless steel guide cannula (C235DCS-5/SPC, Protech International, Boerne, TX) was lowered into the central (stereotaxic coordinates in mm; anteroposterior (AP) +0.7, mediolateral (ML) ±0.4, dorsoventral (DV) −3.3; Paxinos and Franklin, 2001; Dong, 2008) or lateral LS (AP +0.7, ML ±0.6, DV −3.3). Dental cement, formed by mixing Jet denture repair powder (1230P1, Lang Dental Manufacturing, Wheeling, IL) with Jet self-curing acrylic resin liquid (1404CLR, Lang Dental Manufacturing), was applied to secure the cannula pedestal to the top of the skull. The skin at the incision sites was pulled around the implant and sutured around the implant to minimize skull exposure. A mating dummy cannula (C235GS-5/SPC, Protech International, Boerne, TX) was inserted in the guide cannula and secured with a dust cap (303DC/1, Protech International) while the animal recovered in its homecage for at least two weeks.

### 2.2 Intra-LS infusion

Animals were habituated to handling for ten consecutive days (5 min per day). An internal cannula with a 0.25 mm projection at the tip of the guide cannula (C235IS-5/SPC, Protech International) was inserted into the guide cannula. Thin-walled PE50 tubing (C232CT, Protech International) was fitted onto each injector, and the infusate (1 μl total volume per side) was delivered over 4 min and then allowed to penetrate the target region for 2 min before the tubing was removed and attached to the other side of the bilateral cannula. MCH (1 μg; 4025037, Bachem, Torrence, CA), or the MCHR1 antagonist TC-MCH 7c (1 μg; 4365, Tocris, Toronto, Canada) was delivered via a sterile vehicle of artificial cerebrospinal fluid (aCSF) comprising (in mM) 148 NaCl, 3 KCl, 1.4 CaCl_2_, 0.8 MgCl_2_, 0.8 Na_2_HPO_4_, 0.2 NaH_2_PO_4_ prepared as described (ALZET Osmotic Pumps). In experiments where MCH was applied following MCHR1 antagonist pretreatment, TC-MCH 7c (0.5 μg) was infused first and a solution comprising 1 μg MCH and 0.5 µg TC-MCH 7c was infused 5 min later.

### 2.3 Food intake

A chow or 60% high dextrose food pellet was presented on the floor of the homecage. The weight of the food remaining was determined every hour for four hours. Infusions began at ZT 1–3 immediately prior to the start of the 4-h feeding period. Ramekins with regular chow and 60% high dextrose diet were placed on the floor of the homecage overnight and intake of each diet was measured to determine food preference.

### 2.4 Locomotor activity

Micromax infrared beam-break system (Omnitech, Columbus, OH) was used to determine homecage ambulatory activity by recording the total number of beam breaks over four hours following infusion (ZT1–3).

### 2.5 Open field movement

Mice were habituated to an opaque open field arena (45 x 45 x 45 cm) at least three times prior to testing. On the day of testing, sated or fasted mice where food was removed from their homecage for 6 h or 16 h, respectively were habituated to the testing room for 1 h (135 lux). Infusions began at ZT6–11 in the sated condition and ZT1–4 in the fasted condition 30 min prior to testing. Mice were then placed into an opaque open field box (300 lux) with a piece of 60% high dextrose diet affixed to the middle of the arena, and the mice were allowed to explore the arena for 20 min. Movement in the open field arena was recorded with an overhead camera (Hue HD camera, Hue, London, UK) and tracked using ANY-maze (Stoelting Co, Wood Dale, IL). The open field arena was divided into the outside perimeter and center area (22.5 x 22.5 cm). The amount of time the mouse spent in each zone was based on tracking the center-point of the mouse. Plots of animal tracking were generated in ANY-maze and exported as PNG files.

### 2.6 Implant site validation

At the end of the study, mice received an infusion of 0.5% Evans Blue dye in saline (1 µl; MilliporeSigma, Burlington, MA) and were euthanized 30 min later. Mice were anesthetized with chloral hydrate (700 mg/kg; MilliporeSigma) and monitored until reaching a deep anesthetic plane, they were then transcardially perfused with cold (4 °C) saline (0.9% NaCl), followed by fixation with 10% formalin (VWR, Radnor, PA). The brain was extracted from the skull and post-fixed in 10% formalin (VWR, Radnor, PA) for at least 24 h. The brains were cryoprotected in phosphate buffered saline (PBS) containing 20% sucrose and 0.05% sodium azide (24 h, 4 °C) and sliced into five series of 30 µm coronal sections using a freezing microtome (Spencer Lens Co., Buffalo, NY). Brightfield images of LS-containing brain slices were acquired using a Nikon Ti2-E inverted microscope (Nikon Instruments Inc., Mississauga, Canada) and processed using NIS-Elements Imaging Software (Nikon). Cases where the bilateral cannula landed outside the boundaries of the LS, as defined by the Allen Reference Atlas (Dong, 2008), were excluded from analysis.

### 2.7 Experimental design and statistical analyses

Experiments were conducted using a within-group design where mice were tested in the paradigm two to four times, once for each drug condition, in a counterbalanced design. The first hour of food intake was analyzed using a two-tailed, paired *t*-test. Cumulative food intake, locomotor activity, and open field movement were compared by two-way repeated measures ANOVA with Bonferroni post-hoc testing. A three-way ANOVA was used to compare responses between male and female mice over time in their homecage or in the open field. In some cases, a mixed-effect model was used if data points were missing. Results were considered statistically significant at *p* < 0.05. All data graphs were generated using Prism 9 (GraphPad Software, San Diego, CA). Manuscript figures were assembled in Adobe Illustrator 2024 (Adobe Inc., San Jose, CA).

## 3. Results

### 3.1 LS MCH infusion increased chow intake in male and female mice

To determine if the LS mediated the orexigenic actions of MCH, we bilaterally infused MCH (1 µg) into the center of the rostral LS (LSr; **Figure 1A**) of sated male and female mice and measured chow intake over a 4-h period (**Figure 1A**). There was an increase in chow intake within the first hour of MCH infusion (vehicle: 0.06 ± 0.01 g; MCH: 0.11 ± 0.01 g; *t*(10) = 2.93, *p* = 0.015) in male mice leading to a main effect of MCH that more than doubled cumulative intake over 4 h (F(1, 10) = 13.74; *p* = 0.004) and reflected a significant interaction of MCH infusion on chow intake over time (F(4, 40) = 11.46; p < 0.0001; **Figure 1B**). In female mice, MCH infusion into the LS did not significantly increase chow intake in the first hour (vehicle: 0.09 ± 0.03 g; MCH: 0.19 ± 0.07 g; *t*(7) = 1.46, *p* = 0.189), and while we did not detect a significant main effect of MCH (F(1, 7) = 4.75; *p* = 0.066), there was a significant interaction of MCH over time, as MCH increased chow feeding over 4 h (F(4, 28) = 5.49; *p* = 0.002; **Figure 1C**). The orexigenic effect of MCH was comparable between male and female mice, as there was no main effect of sex (F(1, 85) = 2.27; *p* = 0.136) and no interaction between sex and MCH infusion (F(1, 85) = 0.31; *p* = 0.581).

**Figure 1.**
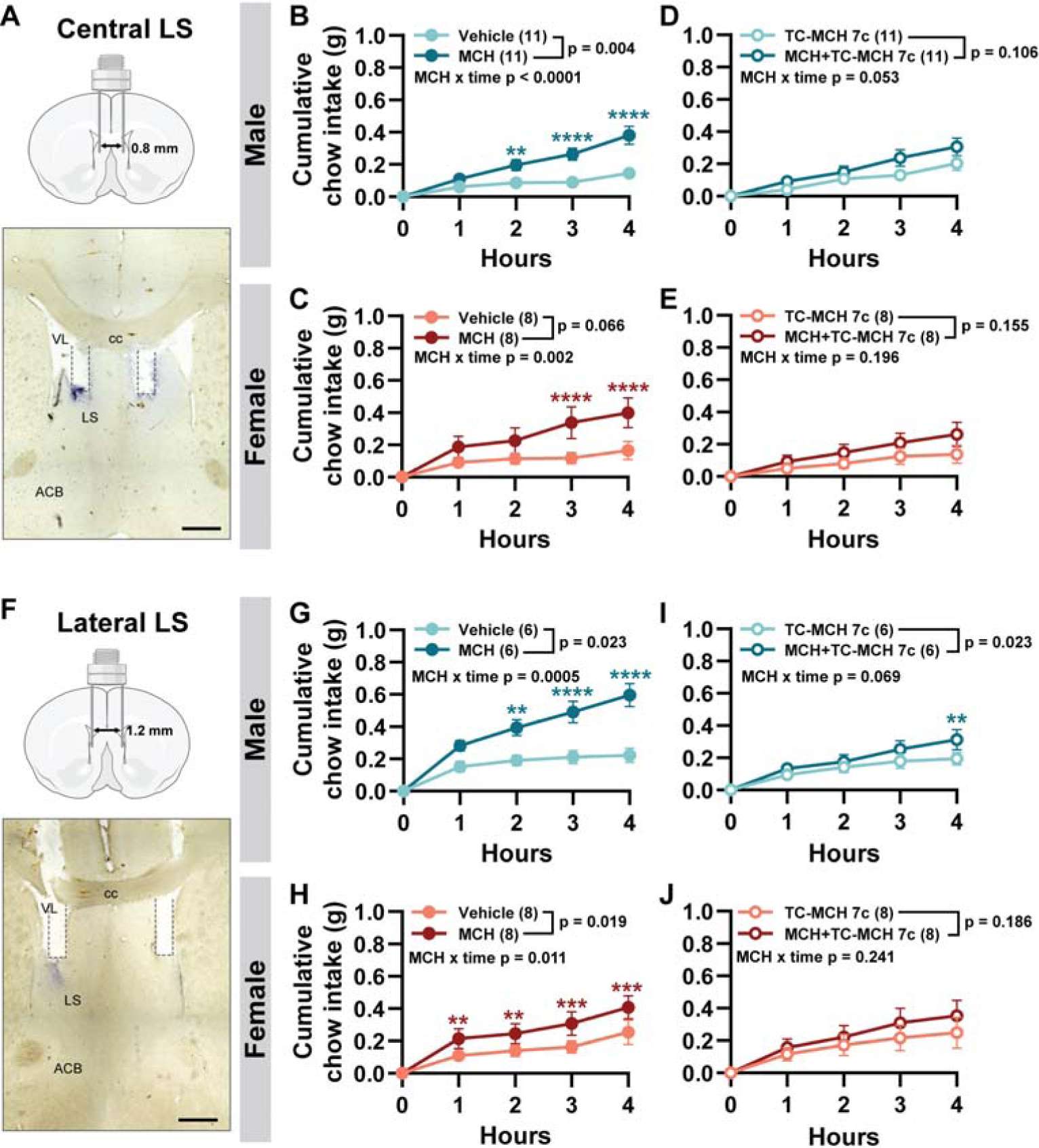
Intra-LS MCH infusion increased chow feeding in male and female mice. Schematic of central LS infusions with bilateral cannulas implanted within 0.4 mm from the midline on each side (*top*). Representative brightfield photomicrograph of infusion spread following dye infusion (*bottom*; **A**). Cumulative chow intake of aCSF vehicle (1 µl) and MCH (1 µg/µl; **B**, **C**) or TC-MCH 7c (1 µg/µl) and MCH in the presence of TC-MCH 7c (1 µg/µl; **D**, **E**) into each side of the central LS of male and female mice. Schematic of lateral LS infusions with bilateral cannulas implanted within 0.6 mm from the midline on each side (*top*). Representative infusion spread following dye infusion (*bottom*; **F**). Cumulative chow intake of aCSF vehicle and MCH (**G**, **H**), or TC-MCH 7c (1 µg/µl) and MCH with TC-MCH 7c (1 µg/µl; **I**, **J**) into each side of the lateral LS of male and female mice. Scale bar: 500 µm (**A**, **F**). Repeated measures two-way ANOVA with Bonferroni post-test comparisons between drug conditions at each timepoint: **, *p* < 0.01; ***, *p* < 0.001; ****, *p* < 0.0001. ACB, nucleus accumbens; cc, corpus callosum; LS, lateral septum; VL, lateral ventricle.

To determine if this orexigenic response was mediated by MCHR1 activation in the LS, we co-infused MCH with the MCHR1 antagonist TC-MCH 7c (1 µg). MCHR1 antagonism has been reported to suppress feeding (Ito et al., 2010), but TC-MCH 7c infusion into the LS alone had no main effect on feeding in male (F(1, 10) = 0.70; *p* = 0.424) or female mice (F(1, 7) = 0.36; *p* = 0.0567) nor a significant interaction of TC-MCH 7c over time in male (F(4, 40) = 1.62; *p* = 0.187) or female mice (F(4, 28) = 0.62; *p* = 0.653) when compared to vehicle infusion. TC-MCH 7c infusion suppressed MCH-mediated feeding, as MCH infusion in the presence of TC-MCH 7c resulted in a similar amount of cumulative feeding in male (F(1, 10) = 3,15; *p* = 0.11) and female mice (F(1, 7) = 2.54; *p* = 0.155). TC-MCH 7c-mediated suppression was sustained over time, as chow intake remained at a similar level over time in both male (F(4, 40) = 2.56; *p* = 0.053; **Figure 1D**) and female mice (F(4, 28) = 1.63; *p* = 0.196; **Figure 1E**) as when the LS was treated with TC-MCH 7c only.

We recently determined that MCHR1 expression concentrated along the lateral LSr border (Payant et al., 2023), thus we performed targeted infusions toward the laterally-distributed MCHR1-expressing LS cells (**Figure 1F**). MCH infusion into the lateral LSr significantly increased feeding within the first hour in both male (vehicle: 0.15 ± 0.03 g; MCH: 0.28 ± 0.03 g; *t*(5) = 3.24, *p* = 0.023) and female mice (vehicle: 0.11 ± 0.02 g; MCH: 0.22 ± 0.06 g; *t*(7) = 2.48, *p* = 0.042). There was a main effect of MCH infusion on cumulative chow intake over four hours in both male (F(1, 5) = 10.38; *p* = 0.023; **Figure 1G**) and female mice (F(1, 7) = 9.19; *p* = 0.019; **Figure 1H**). MCH infusion into the lateral LS of male mice produced a 2-fold increase of chow intake within 2 h (F(4, 20) = 8.13; *p* = 0.0005; **Figure 1G**) and just 1 h in female mice (F(4, 28) = 3.99; *p* = 0.011; **Figure 1H**). Although MCH infusion doubled chow intake relative to vehicle in both male and female mice, there was a main effect of sex (F(1, 60 = 7.18; *p* = 0.010) and an interaction between sex and MCH infusion, which produced a larger increase in chow consumption in male mice (F(1, 60) = 6.72; *p* = 0.012). Furthermore, while both MCH infusion into the central and lateral LS increased feeding over time, the orexigenic effect of MCH was more prominent when infused in the lateral LS, especially in male mice (F(1, 15) = 6.41; *p* = 0.023). This suggested that discrete targeting of MCHR1-rich regions in the lateral LS was most effective at elucidating the orexigenic effects of MCH.

This orexigenic effect was partially blocked by an MCHR1 antagonist in males resulting in a significant main effect (F(1, 5) = 10.45; *p* = 0.023) but no interaction of MCH with TC-MCH 7c over time (F(4, 20) = 2.58; *p* = 0.069; **Figure 1I**). MCH-mediated feeding was fully blocked in females (F(1, 7) = 2.15; *p* = 0.19) and remained at similar levels as TC-MCH 7c alone (F(4, 28) = 1.46; *p* = 0.241; **Figure 1J**). These results indicated that MCH administration in the LS promoted feeding in both male and in female mice.

### 3.2 LS MCH infusion increased palatable food intake in male and female mice

To determine if MCH also promoted hedonic feeding through the LS, we measured the intake of a palatable, high dextrose diet (HDx) over four hours in sated mice following MCH infusion into the central or lateral LS. Mice preferentially consumed HDx (11.3 ± 1.7 kcal/day) over chow (5.7 ± 1.2 kcal/day) thus confirming the palatability of the HDx.

Both male and female mice displayed robust hyperphagia when given access to HDx relative to chow, but when delivered in the central LS, MCH further increased HDx intake in male mice (F(1, 10) = 5.80; *p* = 0.037) over time (F(4, 40) = 3.05; *p* = 0.028; **Figure 2A**). Male mice consumed more HDx overall (F(1. 85) = 16.88; *p* < 0.0001), but there was no significant interaction between sex and MCH infusion (F(1, 85) = 3.44; *p* = 0.067). The orexigenic effect of MCH was blocked by co-infusion of TC-MCH 7c (F(1, 10) = 3.08; *p* = 0.110) and there was a small decrease in feeding over time compared to TC-MCH 7c alone (F(4, 40) = 2.72; *p* = 0.043; **Figure 2B**). By contrast, MCH infusion into the central LS did not further increase HDx intake in female mice (F(1, 7) = 0.024; *p* = 0.880) nor have an effect over time (F(4, 28) = 0.38; *p* = 0.819; **Figure 2C**), and there were also no differences in HDx intake following the co-infusion of MCH with TC-MCH 7c (F(1, 7) = 0.35; *p* = 0.571; MCH x time: F(4, 28) = 0.43; *p* = 0.785).

**Figure 2.**
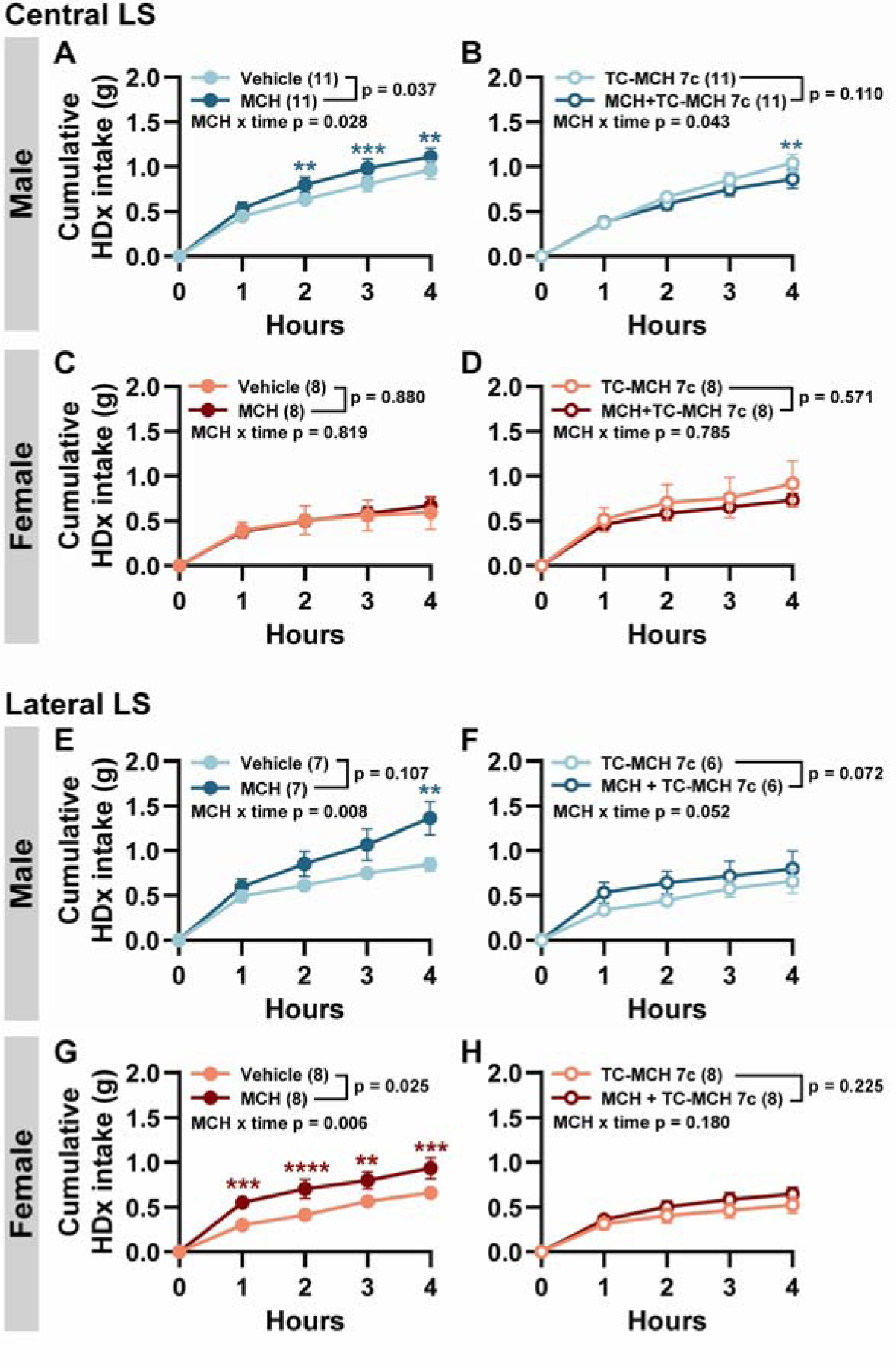
Intra-LS MCH infusion increased palatable diet feeding. Cumulative high dextrose diet (HDx) intake following the infusion of aCSF vehicle (1 µl) and MCH (1 µg/µl) or TC-MCH 7c (1 µg/µl) and MCH with TC-MCH 7c (1 µg/µl) into each side of the central LS of male (**A**, **B**) and female mice (**C**, **D**) or the lateral LS of male (**E**, **F**) and female mice (**G**, **H**). Repeated measures two-way ANOVA with Bonferroni post-test comparisons between drug conditions at each timepoint: **, *p* < 0.01; ***, *p* < 0.001; ****, *p* < 0.0001.

When MCH was infused into the lateral LSr, it increased HDx intake in both sexes, and this orexigenic effect was abolished following TC-MCH 7c co-infusion. In males, there was no main effect of MCH infusion (F(1, 6) = 3.59; *p* = 0.107), but MCH increased HDx intake over time and within 4 h (F(4, 23) = 4.54; *p* = 0.008; **Figure 2E**). This MCH-mediated feeding in the lateral LSr was blocked by TC-MCH 7c co-infusion (F(1, 5) = 5,17; *p* = 0.072; MCH x time: F(4, 20) = 2.83; *p* = 0.052; **Figure 2F**). Interestingly, there was a robust increase in HDx intake following MCH infusion into the lateral LSr of female mice (F(1, 7) = 8.00; *p* = 0.025) that was apparent within an hour of MCH infusion (F(4, 28) = 4.54; *p* = 0.006; **Figure 2G**). This orexigenic MCH effect was blocked by co-infusion of TC-MCH 7c (F(1, 7) = 1.77; *p* = 0.225; MCH x time: F(4, 28) = 1.69; *p* = 0.180; **Figure 2H**). While male mice consumed more of the HDx (F(1, 129) = 18.25; *p* < 0.0001), there was no interaction of sex with MCH infusion (F(1, 129) = 0.11; *p* = 0.741). MCH further increased feeding even when baseline HDx intake was elevated, and MCH-mediated HDx feeding also tended to be greater at the lateral than central LSr of female mice (F(1, 14) = 2.47; *p* = 0.138).

### 3.3 MCH in the lateral LS of male mice increased locomotor activity

While MCH increased food intake when infused in the LS, we did not detect differences in ambulatory or locomotor activity during the feeding period in male mice when MCH was administered alone (F(1, 10) = 0.06; *p* = 0.806; MCH x time: F(3, 30) = 0.29; *p* = 0.830; **Figure 3A**) or in the presence of TC-MCH 7c (F(1, 10) = 0.07; *p* = 0.802; MCH x time: F(3, 30) = 1.35; *p* = 0.278; **Figure 3B**). Similarly, there was no difference in locomotor activity following MCH infusion alone (F(1, 7) = 0.07; *p* = 0.801; MCH x time: F(3, 21) = 1.27; *p* = 0.310; **Figure 3C**) or when MCH was co-infused with TC-MCH 7c (F(1, 7) = 1.16; *p* = 0.317; MCH x time: F(3, 21) = 2.89; *p* = 0.060; **Figure 3D**).

**Figure 3.**
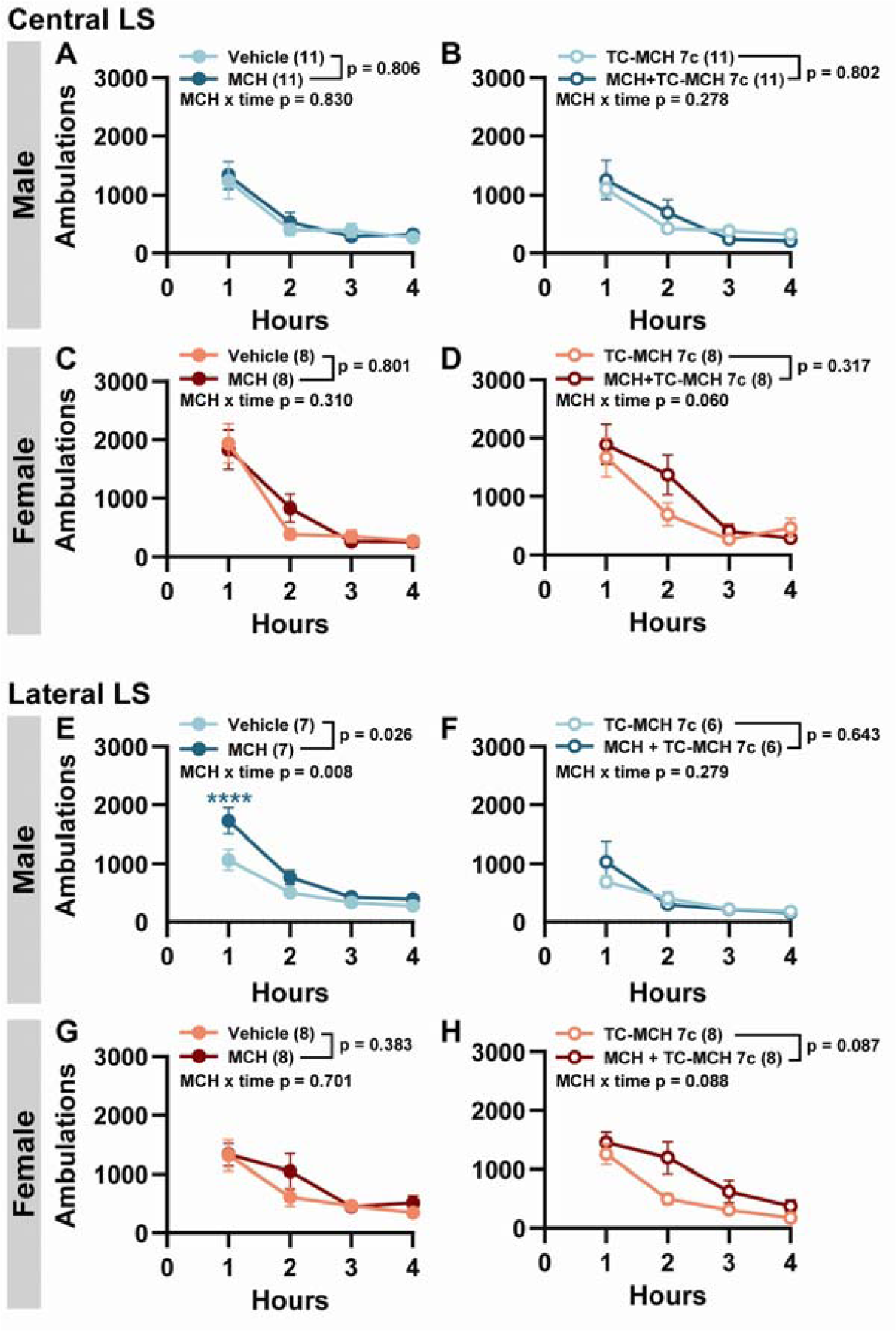
Lateral LS MCH infusion increased locomotor activity in male mice. Total ambulations at each hour following the infusion of aCSF vehicle (1 µl) and MCH (1 µg/µl) or TC-MCH 7c (1 µg/µl) and MCH with TC-MCH 7c (1 µg/µl) into each side of the central LS of male (**A**, **B**) and female mice (**C**, **D**) or the lateral LS of male (**E**, **F**) and female mice (**G**, **H**). Repeated measures two-way ANOVA with Bonferroni post-test comparisons between drug conditions at each timepoint: ****, *p* < 0.0001.

However, when MCH was infused over the lateral LSr, there was an overall increase in ambulatory activity of male mice (F(1, 5) = 9.71; *p* = 0.026). This MCH-mediated hyperactivity in male mice was most prominent within the first hour (F(5, 25) = 4.08; *p* = 0.008; **Figure 3E**) and was abolished by TC-MCH 7c treatment (F(1, 5) = 0.24; *p* = 0.643; MCH x time: F(5, 25) = 1.34; *p* = 0.279; **Figure 3F**). There was no difference in ambulatory activity in female mice when infused with MCH alone (F(1, 7) = 0.86; *p* = 0.383; MCH x time: (F(5, 35) = 0.60; *p* = 0.701; **Figure 3G**) or in the presence of TC-MCH 7c (F(1, 7) = 3.97; *p* = 0.087; MCH x time: F(5, 35) = 2.11; *p* = 0.088; **Figure 3H**). Although the locomotor effect was only present in males, there was no significant interaction between sex and MCH when compared directly (F(1, 72) = 0.255; *p* = 0.615).

### 3.4 LS MCH infusion did not alter food seeking in an anxiogenic environment

As MCH stimulated feeding largely without increasing ambulatory activity, it is possible that MCH-mediate hyperactivity in male mice was attributed to LS roles that regulate anxiogenesis (Bakshi et al., 2007; Xu et al., 2019; Yeates et al., 2022). Furthermore, MCH-mediated feeding may be context-dependent (Subramanian et al., 2023), and while MCH may increase feeding in the homecage, we determined whether MCH would alter food approach behaviours when placed in an anxiogenic environment. We infused MCH into sated mice and placed them into an open field with a HDx food reward placed in the center of the open field to determine if the orexigenic effect of MCH in the LS could overcome their native fear of travelling into the center.

While MCH infusion can drive feeding in sated animals in its homecage, MCH infusion into the central LS did not increase time spent in the center of the open field near the food reward. Female mice spent less time overall exploring the center of the open field (F(1, 10) = 7.0; *p* = 0.025), but there was no main effect of MCH (F(1, 10) = 0.27; *p* = 0.613) or interaction between MCH and sex (F(1, 4) = 0.36; *p* = 0.582; **Figure 4A**). When infused into the lateral LS, male and female mice performed similarly (F(1, 7) = 3.9; *p* = 0.090), but MCH infusion did not alter time spent near the food in the center of the open field(F(1, 7) = 2.2; *p* = 0.182; **Figure 4B**). There was no interaction between sex and MCH (F1, 1) = 0.006; p = 0.952).

**Figure 4.**
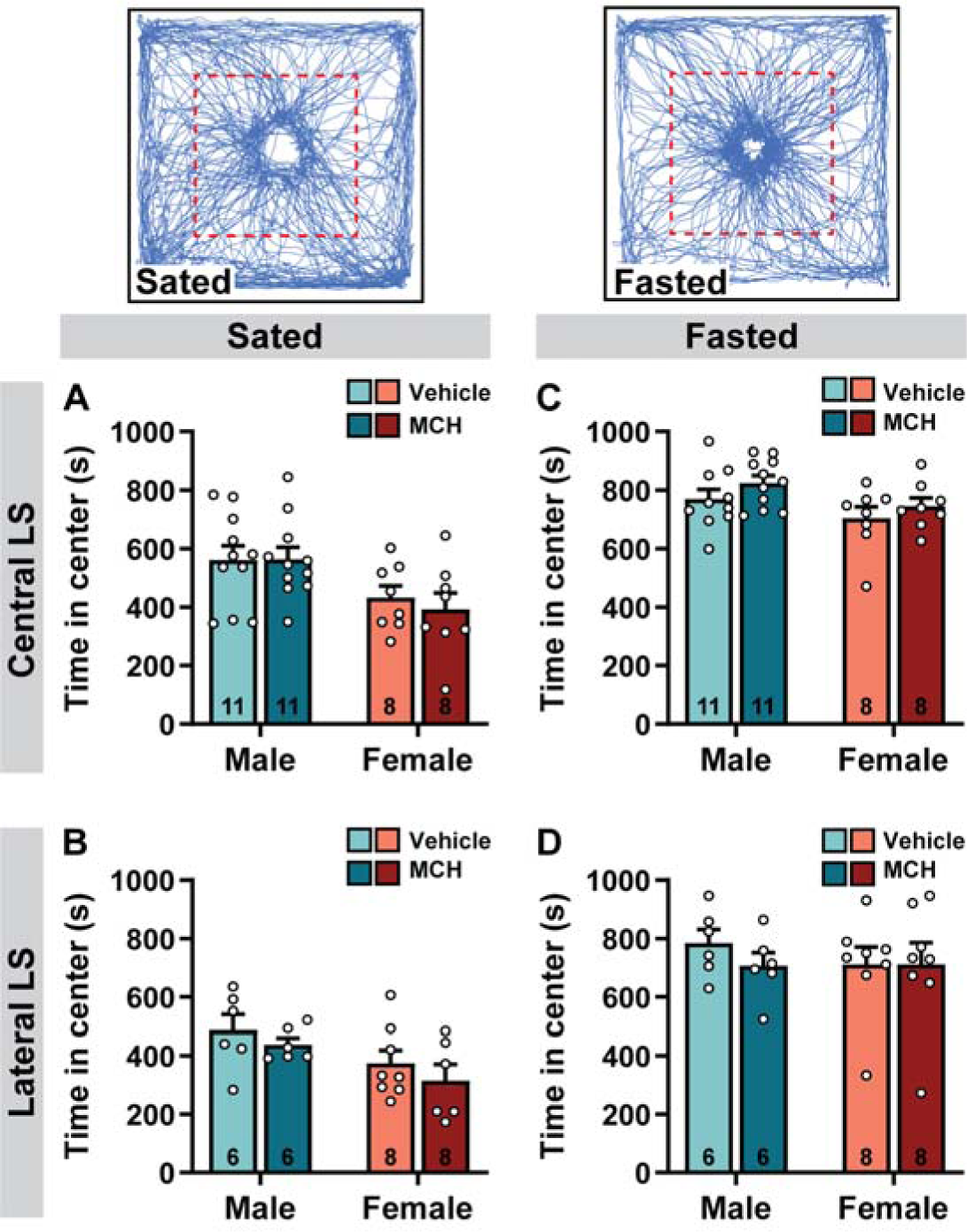
LS MCH infusion did not alter open field food investigation. Representative mouse movement tracks in the center (red outlined area) of an open field in sated and fasted animals (*top*). Time spent in the center of the open field following aCSF vehicle (1 μl) or MCH (1 μg/μl) infusion into each side of the central (A) or lateral LS (B) of sated mice or the central (C) or lateral LS (D) of fasted mice. Significance (*p* < 0.05) was determined using a repeated measures two-way ANOVA.

Hunger is an intrinsic motivator to overcome anxiogenesis and promote food approach behaviours, hence fasted mice will spend more time exploring food placed in the center of an open field (Lockie et al., 2017). We thus determined if MCH infusion in fasted mice promoted food approach in the open field. As anticipated, fasted animals spent more overall time in the center investigating the food reward than sated mice (central LS: (F(1, 17) = 69.65; *p* < 0.0001); lateral LS: (F(1, 12) = 45. 84; *p* < 0.0001). However, MCH did not further increase the amount of time spent in the center when infused into the central (F(1, 10) = 1.10; *p* = 0.324; **Figure 4C**) or lateral LS (F(1, 7) = 0.92; *p* = 0.370; **Figure 4D**). There were no sex differences in the performance of male and female mice in the open field (central LS: F(1, 10) = 4.40; *p* = 0.063); lateral LS: (F(1, 7) = 0.42; *p* = 0.540)), and there was no interaction of sex and MCH infusion (central LS: F(1, 3) = 1.10; *p* = 0.369; lateral LS: (F(1, 3) = 0.97; *p* = 0.398). In sum, the orexigenic effect of MCH in the LS may be restricted to non-stressful environments and insufficient to promote feeding during anxiogenesis.

### 3.5 LS MCH infusion did not alter anxiety-like behaviour

Although MCH infusion did not increase food seeking in an anxiogenic context, the orexigenic effects of MCH may have been masked by MCH-mediated anxiogenesis (Smith et al., 2006; He et al., 2022). We thus determined if MCH infusion in the LS increased anxiety-like behaviour in the open field.

We infused MCH into the central LS and found that while female mice spent less time in the center overall (F(1, 17) = 4.94; *p* = 0.040), there was no main effect of MCH (F(1, 17) = 0.2; *p* = 0.69; **Figure 5A**) or an interaction between sex and MCH (F(1, 17) = 0.89; *p* = 0.359) on time spent in the center of the open field. Consistently, MCH infusion into the lateral LS also did not alter anxiety-like behaviour in the open field in male or female mice (F(1, 12) = 0.1; *p* = 0.744; **Figure 5B**). There was no effect of sex (F(1, 12) = 0.97; *p* = 0.344) or interaction between sex and MCH (F(1, 12) = 0.523; *p* = 0.480). These findings suggested that MCH did not regulate anxiety-like behaviour through the LS and that the lack of feeding response seen in an anxiogenic environment was not masked by an increase in anxiety-like behaviour.

**Figure 5.**
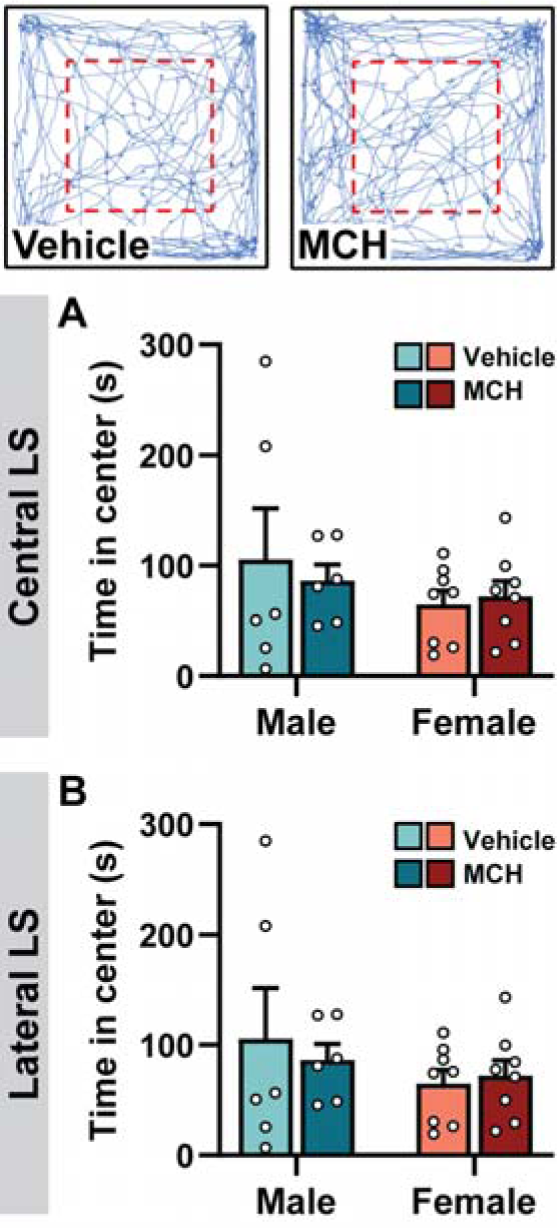
LS MCH infusion did not alter anxiety-like behaviour in the open field. Representative mouse movement tracks (*top*) in the center (red outlined area) of an open field following aCSF vehicle (*left*) or MCH infusion (*right*). Time spent in the center of the open field following vehicle (1 µl) or MCH (1µg/µl) infusion into each side of the central LS (**A**) or lateral LS (**B**). Significant (*p* < 0.05) was determined using a repeated measures two-way ANOVA.

## 4. Discussion

Intracerebroventricular administration of MCH elicited robust feeding effects (Qu et al., 1996; Rossi et al., 1997; Kela et al., 2003), but few brain regions have been identified to support the orexigenic effects of MCH. We showed here that the LS is an important region underlying MCH-mediated feeding. MCH infusion into the LS, especially when directed over the lateral LSr where MCHR1-expressing cells were concentrated (Payant et al., 2023), increased feeding in both male and female mice. However, the orexigenic effects of MCH when the mouse is in its habituated homecage was abolished when placed in an anxiogenic environment, even when the mouse was hungry.

The LS is a novel brain region underlying the orexigenic effect of MCH. Acute MCH infusion into the LS increased consumption of a standard chow or palatable, high-dextrose, diet. MCH induced feeding within the first hour and persisted over four hours which corresponded with the timeline of intracerebroventricular administration of MCH (Qu et al., 1996; Rossi et al., 1997; Kela et al., 2003). MCH infusion into both the central and lateral LS promoted feeding, but consistent with our recent findings showing the concentration of MCHR1 expression towards the lateral LS (Payant et al., 2023), we found that directing MCH towards the lateral LS elicited a more robust orexigenic effect of MCH. The properties of MCH-mediated feeding in the LS are unique from that in the hypothalamus (Rossi et al., 1999; Abbott et al., 2003; Al-Massadi et al., 2019) or nucleus accumbens (Georgescu et al., 2005; Terrill et al., 2020), and it is noteworthy that MCH initiated feeding even during the light cycle when animals were sated. This is in contrast to orexigenic MCH actions in the nucleus accumbens that enhanced chow and sucrose intake in the dark cycle when mice are naturally-feeding but increased only sucrose intake within a 4-hour period in the light cycle (Georgescu et al., 2005; Terrill et al., 2020).

The orexigenic effect of MCH in the LS was uniquely prominent in both male and female mice. Prior studies reported MCH-mediated feeding primarily in males (Terrill et al., 2020), as elevated levels of estrogen in females blunted the orexigenic effect of MCH, and they are restored in ovariectomized females (Messina et al., 2006; Santollo and Eckel, 2008). Estradiol treatment decreases the efficacy of MCH-induced feeding (Messina et al., 2006; Santollo and Eckel, 2008) by reducing the number of MCH cells and downregulating MCHR1 expression in the hypothalamus (Santollo and Eckel, 2013). Although the LS expresses estrogen receptors (Hasunuma et al., 2023), estrogen does not appear to strongly regulate MCH effects in this region. There was a subtle sex difference in MCH-mediated chow consumption whereby MCH infused into the lateral LS of females produced less hyperphagia than in males. However, MCH doubled chow intake in both males and females, therefore the difference in consumption may be related to the higher baseline feeding in males. The distribution and amount of *Mchr1* mRNA or MCHR1 expression is comparable in the LS of male and female mice, and there were no discernable differences in the magnitude of MCH-mediated inhibition in the LS (Payant et al., 2023). The comparable effects of MCH-mediated feeding in the LS of male and female mice were thus consistent with the distribution and functional activation of MCHR1 in the LS.

The orexigenic effect of MCH in LS was context-dependent. MCH-mediated feeding was prominent in the homecage but did not extend to food-seeking in an anxiogenic environment. In the wild, feeding and anxiety are intricately related, as anxiety may suppress feeding but a hungry animal may take risks to feed even in anxiogenic, suboptimal environments. We observed a mild increase in locomotor activity in male mice when MCH was infused into the lateral LS. Since this effect was absent in the other groups exhibiting feeding effects, it appeared to be unrelated to feeding. As such, we hypothesized that the hyperactivity could be related to an anxiolytic or exploratory behavioural outcome. LS inhibition can suppress anxiety-like behaviour (Pesold and Treit, 1996; Wang et al., 2023) and the switch to exploratory behaviour in the face of risk (Zhong et al., 2022). Therefore, we determined if MCH infusion into the LS may increase time spent investigating a food reward placed in the center of an open field. However, in sated or fasted mice, the orexigenic drive produced by MCH did not increase the amount of time mice spent in the center of the open field and this was not due to an opposing increase in anxiety-like behaviour. While MCH in the LS did not mediate the interaction between feeding and anxiety, it may still extend beyond homecage feeding to other facets of feeding-related behaviour. The LS receives strong input from the hippocampus (Risold and Swanson, 1997), which promotes spatial food memory (Decarie-Spain et al., 2022), and LS activation shortens the latency to food approach (Chen et al., 2022). As the infusion of MCH or activation of MCH terminals in the LS enhanced spatial memory by potentiating hippocampal inputs (Liu et al., 2022), future studies may evaluate whether MCH acts in the LS to regulate spatial memory-dependent food-seeking.

The magnitude of the orexigenic effect with intra-LS MCH infusion was larger with chow feeding than palatable diet feeding. This result was unexpected because MCH is thought to reinforce the consumption of calorically dense solutions (Duncan et al., 2005; Karlsson et al., 2012) by increasing their hedonic value (Baird et al., 2006; Lopez et al., 2011) and by recruiting post-oral processes that detect and reinforce the consumption of nutrient-dense substances (Domingos et al., 2013). However, this is often shown through the comparison of sweetened solutions in the presence and absence of calories, for example via sucrose and sucralose, respectively, and therefore differential outcomes could be expected when comparing two nutrient-containing solid diets. Furthermore, as baseline feeding was greatly elevated when given access to HDx, ceiling levels of baseline palatable HDx feeding would mask any orexigenic effects of MCH, such as that seen with standard chow.

Chronic MCHR1 antagonism can suppress feeding (Shearman et al., 2003; Ito et al., 2010; Kawata et al., 2017), but acute MCHR1 antagonism has no effect (Shearman et al., 2003) or may decrease feeding in the dark cycle on feeding (Georgescu et al., 2005). Unsurprisingly, we did not see any reductions in feeding within four hours of LS infusion during the light cycle. We noted that administration of TC-MCH 7c with MCH did not fully block the effects of MCH in males. It is not clear if this is due to the timing or insufficiency of MCHR1 antagonism. We administered TC-MCH 7c a few minutes prior to MCH infusion and this may have been insufficient time for the antagonist to block MCHR1 prior to the co-infusion of TC-MCH 7c with MCH. Additionally, TC-MCH 7c may not fully block the cellular effects of MCH, as TC-MCH 7c pretreatment at LS cells only partly blocked the inhibitory effects of MCH (Liu et al., 2022).

## 5. Conclusion

MCH infusion into the LS promoted hyperphagia in male and female mice and these results implicated the LS as an important structure underlying MCH-mediated behaviour. As prior studies were conducted in rat, cross species differences hindered our ability to assess the relative contribution of the LS to MCH-mediated feeding compared to other brain targets.

However, MCH uniquely initiated feeding via the LS, and as LS inhibition can stimulate feeding (Mitra et al., 2014; Gabriella et al., 2022), and MCH inhibits LS cells (Payant et al., 2023), the MCH system is an emerging regulator of LS-mediated feeding.

## Author contributions

Conceptualization: M.J.C. Methodology: M.A.P, M.J.C. Investigation: A.S, M.A.P, M.J.C. Formal Analysis: A.S, M.A.P. Visualization: M.A.P. Writing – original draft: M.A.P. Writing – review & editing: M.A.P, A.S., M.J.C.

## Data availability

The data from this study are available upon reasonable request.

## Acknowledgements

This work was supported by Natural Sciences and Engineering Research Council of Canada (NSERC) Discovery Grant RGPIN-2024-04133 (MJC), NSERC Postgraduate Doctoral Scholarship (MAP), and NSERC Undergraduate Student Research Award (AS), as well as by the Province of Ontario Graduate Scholarship (MAP).

## References

Abbott CR, Kennedy AR, Wren AM, Rossi M, Murphy KG, Seal LJ, Todd JF, Ghatei MA, Small CJ, Bloom SR (2003) Identification of hypothalamic nuclei involved in the orexigenic effect of melanin-concentrating hormone. Endocrinology 144:3943–3949.

Al-Massadi O et al. (2019) MCH Regulates SIRT1/FoxO1 and Reduces POMC Neuronal Activity to Induce Hyperphagia, Adiposity, and Glucose Intolerance. Diabetes 68:2210–2222.

Baird JP, Rios C, Gray NE, Walsh CE, Fischer SG, Pecora AL (2006) Effects of melanin-concentrating hormone on licking microstructure and brief-access taste responses. Am J Physiol Regul Integr Comp Physiol 291:R1265–1274.

Bakshi VP, Newman SM, Smith-Roe S, Jochman KA, Kalin NH (2007) Stimulation of lateral septum CRF2 receptors promotes anorexia and stress-like behaviors: functional homology to CRF1 receptors in basolateral amygdala. J Neurosci 27:10568–10577.

Chee MJ, Pissios P, Maratos-Flier E (2013) Neurochemical characterization of neurons expressing melanin-concentrating hormone receptor 1 in the mouse hypothalamus. J Comp Neurol 521:2208–2234.

Chee MJ, Arrigoni E, Maratos-Flier E (2015) Melanin-concentrating hormone neurons release glutamate for feedforward inhibition of the lateral septum. J Neurosci 35:3644–3651.

Chen Z, Chen G, Zhong J, Jiang S, Lai S, Xu H, Deng X, Li F, Lu S, Zhou K, Li C, Liu Z, Zhang X, Zhu Y (2022) A circuit from lateral septum neurotensin neurons to tuberal nucleus controls hedonic feeding. Mol Psychiatry 27:4843–4860.

Decarie-Spain L, Liu CM, Lauer LT, Subramanian K, Bashaw AG, Klug ME, Gianatiempo IH, Suarez AN, Noble EE, Donohue KN, Cortella AM, Hahn JD, Davis EA, Kanoski SE (2022) Ventral hippocampus-lateral septum circuitry promotes foraging-related memory. Cell Rep 40:111402.

Dilsiz P, Aklan I, Sayar Atasoy N, Yavuz Y, Filiz G, Koksalar F, Ates T, Oncul M, Coban I, Ates Oz E, Cebecioglu U, Alp MI, Yilmaz B, Atasoy D (2020) MCH Neuron Activity Is Sufficient for Reward and Reinforces Feeding. Neuroendocrinology 110:258–270.

Diniz GB, Battagello DS, Klein MO, Bono BSM, Ferreira JGP, Motta-Teixeira LC, Duarte JCG, Presse F, Nahon JL, Adamantidis A, Chee MJ, Sita LV, Bittencourt JC (2020) Ciliary melanin-concentrating hormone receptor 1 (MCHR1) is widely distributed in the murine CNS in a sex-independent manner. J Neurosci Res 98:2045–2071.

Domingos AI, Sordillo A, Dietrich MO, Liu ZW, Tellez LA, Vaynshteyn J, Ferreira JG, Ekstrand MI, Horvath TL, de Araujo IE, Friedman JM (2013) Hypothalamic melanin concentrating hormone neurons communicate the nutrient value of sugar. Elife 2:e01462.

Dong H (2008) The Allen reference atlas: A digital color brain atlas of the C57BL/6J male mouse.. Hoboken, NJ: John Wiley & Sons.

Duncan EA, Proulx K, Woods SC (2005) Central administration of melanin-concentrating hormone increases alcohol and sucrose/quinine intake in rats. Alcohol Clin Exp Res 29:958–964.

Elliott JC, Harrold JA, Brodin P, Enquist K, Backman A, Bystrom M, Lindgren K, King P, Williams G (2004) Increases in melanin-concentrating hormone and MCH receptor levels in the hypothalamus of dietary-obese rats. Brain Res Mol Brain Res 128:150–159.

Gabriella I, Tseng A, Sanchez KO, Shah H, Stanley BG (2022) Stimulation of GABA Receptors in the Lateral Septum Rapidly Elicits Food Intake and Mediates Natural Feeding. Brain Sci 12.

Georgescu D, Sears RM, Hommel JD, Barrot M, Bolanos CA, Marsh DJ, Bednarek MA, Bibb JA, Maratos-Flier E, Nestler EJ, DiLeone RJ (2005) The hypothalamic neuropeptide melanin-concentrating hormone acts in the nucleus accumbens to modulate feeding behavior and forced-swim performance. J Neurosci 25:2933–2940.

Guesdon B, Paradis E, Samson P, Richard D (2009) Effects of intracerebroventricular and intra-accumbens melanin-concentrating hormone agonism on food intake and energy expenditure. Am J Physiol Regul Integr Comp Physiol 296:R469–475.

Hasunuma K, Murakawa T, Takenawa S, Mitsui K, Hatsukano T, Sano K, Nakata M, Ogawa S (2023) Estrogen Receptor beta in the Lateral Septum Mediates Estrogen Regulation of Social Anxiety-like Behavior in Male Mice. Neuroscience.

He X, Li Y, Zhang N, Huang J, Ming X, Guo R, Hu Y, Ji P, Guo F (2022) Melanin-concentrating hormone promotes anxiety and intestinal dysfunction via basolateral amygdala in mice. Front Pharmacol 13:906057.

Ito M, Ishihara A, Gomori A, Matsushita H, Ito M, Metzger JM, Marsh DJ, Haga Y, Iwaasa H, Tokita S, Takenaga N, Sato N, MacNeil DJ, Moriya M, Kanatani A (2010) Mechanism of the anti-obesity effects induced by a novel melanin-concentrating hormone 1-receptor antagonist in mice. Br J Pharmacol 159:374–383.

Izawa S, Chowdhury S, Miyazaki T, Mukai Y, Ono D, Inoue R, Ohmura Y, Mizoguchi H, Kimura K, Yoshioka M, Terao A, Kilduff TS, Yamanaka A (2019) REM sleep-active MCH neurons are involved in forgetting hippocampus-dependent memories. Science 365:1308–1313.

Karlsson C, Zook M, Ciccocioppo R, Gehlert DR, Thorsell A, Heilig M, Cippitelli A (2012) Melanin-concentrating hormone receptor 1 (MCH1-R) antagonism: reduced appetite for calories and suppression of addictive-like behaviors. Pharmacol Biochem Behav 102:400–406.

Kawata Y, Okuda S, Hotta N, Igawa H, Takahashi M, Ikoma M, Kasai S, Ando A, Satomi Y, Nishida M, Nakayama M, Yamamoto S, Nagisa Y, Takekawa S (2017) A novel and selective melanin-concentrating hormone receptor 1 antagonist ameliorates obesity and hepatic steatosis in diet-induced obese rodent models. Eur J Pharmacol 796:45–53.

Kela J, Salmi P, Rimondini-Giorgini R, Heilig M, Wahlestedt C (2003) Behavioural analysis of melanin-concentrating hormone in rats: evidence for orexigenic and anxiolytic properties. Regul Pept 114:109–114.

Lee J, Raycraft L, Johnson AW (2021) The dynamic regulation of appetitive behavior through lateral hypothalamic orexin and melanin concentrating hormone expressing cells. Physiol Behav 229:113234.

Lembo PM, Grazzini E, Cao J, Hubatsch DA, Pelletier M, Hoffert C, St-Onge S, Pou C, Labrecque J, Groblewski T, O’Donnell D, Payza K, Ahmad S, Walker P (1999) The receptor for the orexigenic peptide melanin-concentrating hormone is a G-protein-coupled receptor. Nat Cell Biol 1:267–271.

Liu JJ, Tsien RW, Pang ZP (2022) Hypothalamic melanin-concentrating hormone regulates hippocampus-dorsolateral septum activity. Nat Neurosci 25:61–71.

Lockie SH, McAuley CV, Rawlinson S, Guiney N, Andrews ZB (2017) Food Seeking in a Risky Environment: A Method for Evaluating Risk and Reward Value in Food Seeking and Consumption in Mice. Front Neurosci 11:24.

Lopez CA, Guesdon B, Baraboi ED, Roffarello BM, Hetu M, Richard D (2011) Involvement of the opioid system in the orexigenic and hedonic effects of melanin-concentrating hormone. Am J Physiol Regul Integr Comp Physiol 301:R1105–1111.

Messina MM, Boersma G, Overton JM, Eckel LA (2006) Estradiol decreases the orexigenic effect of melanin-concentrating hormone in ovariectomized rats. Physiol Behav 88:523–528.

Mitra A, Lenglos C, Timofeeva E (2014) Activation of GABAA and GABAB receptors in the lateral septum increases sucrose intake by differential stimulation of sucrose licking activity. Behav Brain Res 273:82–88.

Mul JD, la Fleur SE, Toonen PW, Afrasiab-Middelman A, Binnekade R, Schetters D, Verheij MM, Sears RM, Homberg JR, Schoffelmeer AN, Adan RA, DiLeone RJ, De Vries TJ, Cuppen E (2011) Chronic loss of melanin-concentrating hormone affects motivational aspects of feeding in the rat. PLoS One 6:e19600.

Noble EE, Hahn JD, Konanur VR, Hsu TM, Page SJ, Cortella AM, Liu CM, Song MY, Suarez AN, Szujewski CC, Rider D, Clarke JE, Darvas M, Appleyard SM, Kanoski SE (2018) Control of Feeding Behavior by Cerebral Ventricular Volume Transmission of Melanin-Concentrating Hormone. Cell Metab 28:55–68 e57.

Paxinos G, Franklin K (2001) The Mouse Brain in Stereotaxic Coordinates. San Diego, CA: Academic Press.

Payant MA, Spencer CD, Chee MJ (2023) Inhibitory actions of melanin-concentrating hormone in the lateral septum. bioRxiv. doi: 10.1101/2023.10.21.562777

Pesold C, Treit D (1996) The neuroanatomical specificity of the anxiolytic effects of intra-septal infusions of midazolam. Brain Res 710:161–168.

Pissios P, Frank L, Kennedy AR, Porter DR, Marino FE, Liu FF, Pothos EN, Maratos-Flier E (2008) Dysregulation of the mesolimbic dopamine system and reward in MCH-/- mice. Biol Psychiatry 64:184–191.

Qu D, Ludwig DS, Gammeltoft S, Piper M, Pelleymounter MA, Cullen MJ, Mathes WF, Przypek R, Kanarek R, Maratos-Flier E (1996) A role for melanin-concentrating hormone in the central regulation of feeding behaviour. Nature 380:243–247.

Risold PY, Swanson LW (1997) Connections of the rat lateral septal complex. Brain Res Brain Res Rev 24:115–195.

Romero-Pico A, Sanchez-Rebordelo E, Imbernon M, Gonzalez-Touceda D, Folgueira C, Senra A, Ferno J, Blouet C, Cabrera R, van Gestel M, Adan RA, Lopez M, Maldonado R, Nogueiras R, Dieguez C (2018) Melanin-Concentrating Hormone acts through hypothalamic kappa opioid system and p70S6K to stimulate acute food intake. Neuropharmacology 130:62–70.

Rossi M, Choi SJ, O’Shea D, Miyoshi T, Ghatei MA, Bloom SR (1997) Melanin-concentrating hormone acutely stimulates feeding, but chronic administration has no effect on body weight. Endocrinology 138:351–355.

Rossi M, Beak SA, Choi SJ, Small CJ, Morgan DG, Ghatei MA, Smith DM, Bloom SR (1999) Investigation of the feeding effects of melanin concentrating hormone on food intake--action independent of galanin and the melanocortin receptors. Brain Res 846:164–170.

Saito Y, Cheng M, Leslie FM, Civelli O (2001) Expression of the melanin-concentrating hormone (MCH) receptor mRNA in the rat brain. J Comp Neurol 435:26–40.

Santollo J, Eckel LA (2008) The orexigenic effect of melanin-concentrating hormone (MCH) is influenced by sex and stage of the estrous cycle. Physiol Behav 93:842–850.

Santollo J, Eckel LA (2013) Oestradiol decreases melanin-concentrating hormone (MCH) and MCH receptor expression in the hypothalamus of female rats. J Neuroendocrinol 25:570–579.

Shearman LP, Camacho RE, Sloan Stribling D, Zhou D, Bednarek MA, Hreniuk DL, Feighner SD, Tan CP, Howard AD, Van der Ploeg LH, MacIntyre DE, Hickey GJ, Strack AM (2003) Chronic MCH-1 receptor modulation alters appetite, body weight and adiposity in rats. Eur J Pharmacol 475:37–47.

Sherwood A, Holland PC, Adamantidis A, Johnson AW (2015) Deletion of Melanin Concentrating Hormone Receptor-1 disrupts overeating in the presence of food cues. Physiol Behav 152:402–407.

Smith DG, Davis RJ, Rorick-Kehn L, Morin M, Witkin JM, McKinzie DL, Nomikos GG, Gehlert DR (2006) Melanin-concentrating hormone-1 receptor modulates neuroendocrine, behavioral, and corticolimbic neurochemical stress responses in mice. Neuropsychopharmacology 31:1135–1145.

Subramanian KS, Lauer LT, Hayes AMR, Decarie-Spain L, McBurnett K, Nourbash AC, Donohue KN, Kao AE, Bashaw AG, Burdakov D, Noble EE, Schier LA, Kanoski SE (2023) Hypothalamic melanin-concentrating hormone neurons integrate food-motivated appetitive and consummatory processes in rats. Nat Commun 14:1755.

Terrill SJ, Subramanian KS, Lan R, Liu CM, Cortella AM, Noble EE, Kanoski SE (2020) Nucleus accumbens melanin-concentrating hormone signaling promotes feeding in a sex-specific manner. Neuropharmacology 178:108270.

Wang D, Pan X, Zhou Y, Wu Z, Ren K, Liu H, Huang C, Yu Y, He T, Zhang X, Yang L, Zhang H, Han MH, Liu C, Cao JL, Yang C (2023) Lateral septum-lateral hypothalamus circuit dysfunction in comorbid pain and anxiety. Mol Psychiatry 28:1090–1100.

Xu Y, Lu Y, Cassidy RM, Mangieri LR, Zhu C, Huang X, Jiang Z, Justice NJ, Xu Y, Arenkiel BR, Tong Q (2019) Identification of a neurocircuit underlying regulation of feeding by stress-related emotional responses. Nat Commun 10:3446.

Yeates DCM, Leavitt D, Sujanthan S, Khan N, Alushaj D, Lee ACH, Ito R (2022) Parallel ventral hippocampus-lateral septum pathways differentially regulate approach-avoidance conflict. Nat Commun 13:3349.

Zhong C, Wang L, Cao Y, Sun C, Huang J, Wang X, Pan S, He S, Huang K, Lu Z, Xu F, Lu Y, Wang L (2022) A neural circuit from the dorsal CA3 to the dorsomedial hypothalamus mediates balance between risk exploration and defense. Cell Rep 41:111570.

